# Profiling the transcallosal response of rat motor cortex evoked by contralateral optogenetic stimulation of glutamatergic cortical neurons

**DOI:** 10.1101/2021.04.15.439619

**Authors:** Christian Stald Skoven, Leo Tomasevic, Duda Kvitsiani, Bente Pakkenberg, Tim Bjørn Dyrby, Hartwig Roman Siebner

## Abstract

**Background:** Efficient interhemispheric integration of neural activity between left and right primary motor cortex (M1) is critical for inter-limb motor control.

**Objective:** We employed optogenetic stimulation to establish a framework for probing transcallosal M1-M1 interactions in rats.

**Methods:** In male rats, we optogenetically stimulated glutamatergic neurons in right M1 and recorded the transcallosally evoked potential with chronically implanted electrodes in contralateral left M1 during dexmedetomidine anesthesia. We systematically varied the stimulation intensity and duration to characterize the relationship between stimulation parameters in right M1 and the characteristics of the evoked intracortical potentials in left M1.

**Results:** Optogenetic stimulation of right M1 consistently evoked a transcallosal response in left M1 with a consistent negative peak (N1) that sometimes was preceded by a smaller positive peak (P1). Higher stimulation intensity or longer stimulation duration gradually increased N1 amplitude and reduced N1 variability across trials. Median N1 latencies remained stable, once stimulation elicited a reliable N1 peak and did not display a systematic shortening with increasing stimulation intensity or duration.

**Conclusions:** Optogenetically stimulated glutamatergic neurons in M1 can reliably evoke a transcallosal response in anesthetized rats and can be used to characterize the relationship between “stimulation dose” and “response magnitude” (i.e., the gain function) of transcallosal M1-to-M1 glutamatergic connections. Detailed knowledge of the stimulus-response relationship is needed to optimize the efficacy of optogenetic stimulation. Since transcallosal M1-M1 interactions can also be probed non-invasively with transcranial magnetic stimulation in humans, our optogenetic stimulation approach bears translational potential for studying how unilateral M1 stimulation can induce interhemispheric plasticity.

## Introduction

The corpus callosum (CC) connects homologous cortical sites in the right and left hemisphere and therefore is a critical structure for inter-hemispheric integration in the mammalian brain [1,2]. This also applies to the premotor and primary sensorimotor cortices. In primates, transcallosal fibers provide an important pathway through which somatosensory information and motor commands from the right and left limbs are integrated with substantial differences in interhemispheric connectivity among cortical areas [3,4]. Early neurophysiological studies applied cortical electrical stimulation of the cortex in one hemisphere of animals and recorded the “callosal potentials” that were elicited in the homologous part of the opposite hemisphere [5]. Severing the CC at the midline completely abolished the electrically evoked potentials [5]. Augmenting the intensity of electrical stimulation, the recorded peak amplitude in the opposite hemisphere gradually increased without a change in the latency of the peak [6]. The peak appeared sharper when the CC was stimulated directly [6], indicating a more synchronized response.

In humans, transcallosal conduction between the two motor hand areas can be probed non-invasively with dual-coil transcranial magnetic stimulation (TMS) [7–9]. A conditioning stimulus applied to the motor hand area of one hemisphere inhibits the excitability of the contralateral motor hand area, which is probed with a test stimulus given to the contralateral motor cortex. The latency of IHI reflects the transcallosal conduction time [7]. The strength of interhemispheric inhibition (IHI) dynamically changes depending on the motor context and may be altered by interventional transcranial stimulation [10–13] or by brain diseases [14–18].

The advent of opto- [19,20] and pharmacogenetic [21,22] tools to selectively stimulate a distinct class of brain cells has massively expanded the potential of studying the CC and its function in animals - both *in vitro* [23] and *in vivo* [24–27]. Optogenetic stimulation of the transcallosal somatosensory or motor projections provides a powerful tool to study the causal significance of the corpus callosum in interhemispheric sensorimotor integration. Optogenetic stimulation of the transcallosal somatosensory or motor projections provides a precise interventional tool to study directional functional connectivity from one sensorimotor cortex to its hemispheric counterpart. But appropriate use of optogenetic interventions requires detailed knowledge about the stimulus-response relationship between the stimulation variables and the evoked neuronal response in the contralateral hemisphere. Recent optogenetic studies in rodents and nonhuman primates emphasize that the stimulation induced neuronal activity in the brain, such as the size of the evoked local field potential (LFP) or the change in axonal firing rates, depends on the stimulation variables, such as the stimulation intensity applied or the duration of laser stimulation [25,27,28]. Although the impact of key variables might have been investigated systematically beforehand in many optogenetic stimulation studies, it often remains unclear why a specific set of stimulation variables was used for optogenetic stimulation.

In this study, we combined optogenetic stimulation with intracortical electrophysiological recordings to characterize functional transcallosal motor-to-motor interactions in anesthetized rats. To establish a robust experimental framework for future studies, we conducted an experiment to identify the optimal stimulation settings for optogenetic stimulation of glutamatergic neurons in the motor cortex (M1) and their transcallosal projections to the contralateral homologous M1. We hypothesized that the magnitude of the optogenetically evoked intracortical response would gradually increase with stimulation duration and intensity.

## Methods

All animal procedures were conducted in accordance with the Danish and EU legislation and were approved by The Animal Experiments Inspectorate (2016-15-0201-00868). Figure 1 illustrates our experimental approach. In young male rats (NTac:SD-M), we stimulated glutamatergic neurons in the right M1 with a chronically implanted optic fiber and recorded the transcallosal cortical response with an intracortical electrode implanted in the contralateral M1. We systematically investigated the relationship between two key variables of optogenetic stimulation, namely stimulation duration and intensity, and two electrophysiological measures, the latency and amplitude of the transcallosal evoked LFP.

**Figure 1:**
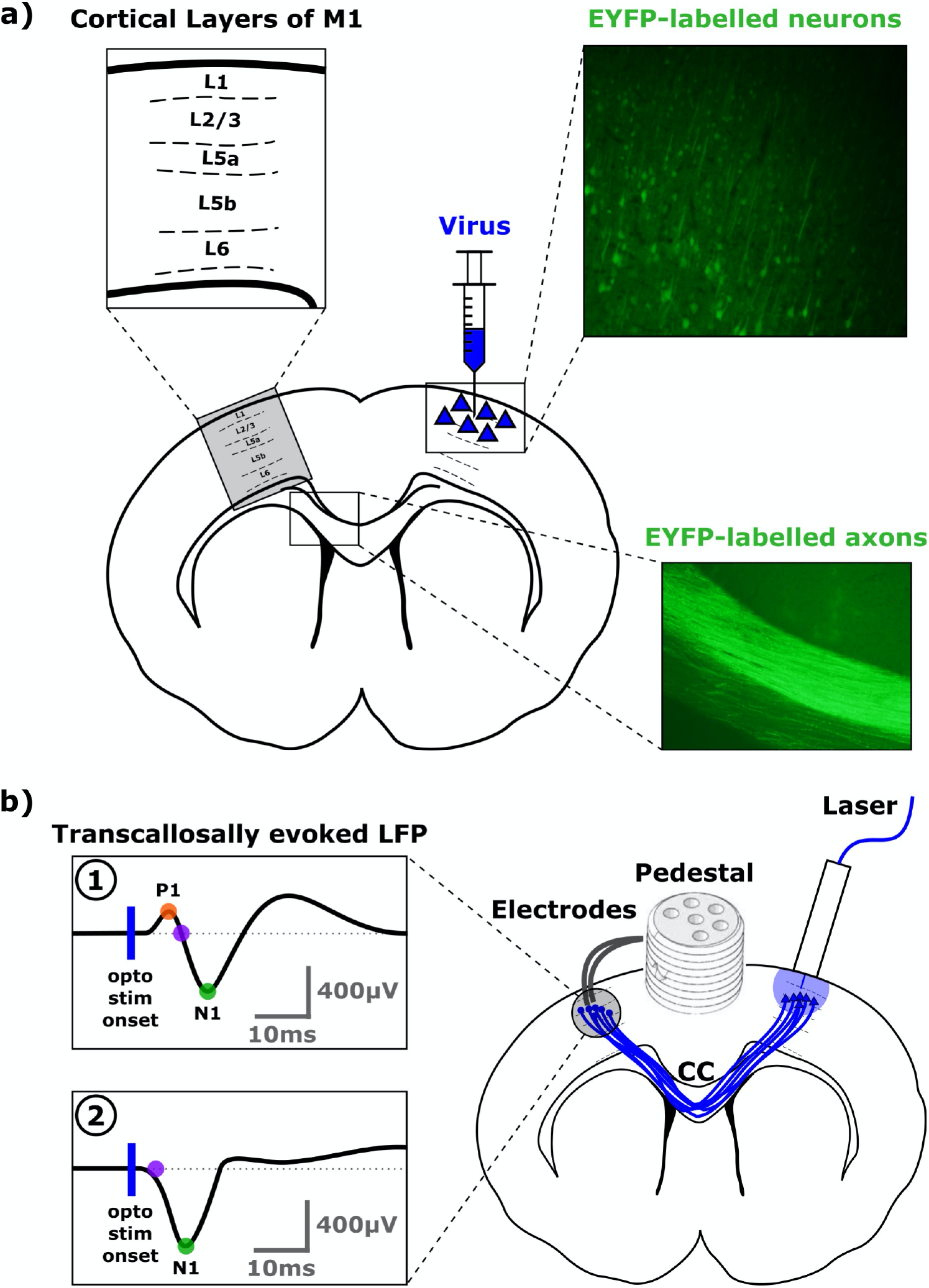
Synopsis of the optogenetic and electrophysiological experimental procedures. The anatomical drawings depict coronal sections (adapted from [30]) of the rat brain at +1.00mm anterior to bregma according to [30]. The black filled areas indicate the ventricles, while the white central area in the brain corresponds to cerebral white matter, including corpus callosum (CC). **Panel a)** The viral optogenetic construct was injected in the upper layers of right M1. As a result, Channelrhodopsin2 (ChR2) and Enhanced Yellow Fluorescent Protein (EYFP) were expressed in the neurons at the injection site and along those axonal projections, projecting to the contralateral hemisphere via the corpus callosum (CC). Bright green color in the fluorescence microscopy inserts corresponds to EYFP expressed alongside ChR2. **Panel b)** An optic fiber was implanted in right M1 for optogenetic excitation of the transcallosal glutamatergic projection fibers from right to left M1 (depicted as blue lines). A stereo-electrode was implanted in contralateral left M1 to record the transcallosally evoked Local Field Potentials (LFP) after optogenetic stimulation. We recorded two morphologies of transcallosally evoked LFP responses which are illustrated in the left lower part of the figure. **(1)** The majority of LFP responses showed an initial positive deflection followed by a second negative deflection. The first positive peak (P1) is marked by an orange dot. The first negative peak (N1) is marked by a green dot. The onset latency of N1 (purple dot) was interpolated to the baseline, based on the slope around mid-maximum of the N1 peak. **(2)** Some LFP responses lacked an initial positive component and started directly with a negative deflection (purple dot). The vertical blue line depicts the onset of optogenetic stimulation.

### Construction and calibration of the fiber implant

The fiber implants were constructed as follows: a multimode optical fiber (Ø = 55 μm, 3 mm protruding; FVP050055065, Polymicro Technologies LLC, CM Scientific) fixed in a ceramic ferrule (L = 6.4 mm, ID: 127 μm, OD: 1.25 mm; MM-FER2007C-1260P, PFP) with cyanoacrylate glue (Loctite Universal). After hardening, every fiber implant was successively polished (30 μm, 6 μm and 3 μm; ThorLabs: LF30D, LF6D, LF3D). Before implantation, the stimulation intensity was calibrated for each optic fiber implant [29]. The optic fiber implants were connected (via ADAL1 or ADAL3, Thorlabs) to a custom-made fiber patch cable (5 m, Ø = 50 μm, Thorlabs), which in turn was connected (via ADAFC1) to the fiber-coupled (1 m, Ø = 50 μm, FC/PC) laser source (LaserMate BML447-150FLAM5F, 447 nm, 500 mW). The laser was turned on and allowed to stabilize for approximately 15 minutes [29]. Stimulation intensity was measured at a fixed distance and position, by a custom-made 3D-printed (with InnoFil Black Pro1 on Ultimaker 2+ extended) holder (https://git.drcmr.dk/cskoven/lab), in relation to the sensor (S121C, ThorLabs) connected to the PowerMeter (PM100A, ThorLabs).

The analog dial setting on the laser source was then noted for the different stimulation intensities [0.25, 0.50, 1.00, 2.00, 5.00, 10.00] mW (from ~105 mW/mm^2^ to ~4209 mW/mm^2^) for each fiber implant.

### Surgical procedures

21 young male rats (NTac:SD-M; 3weeks old) were received and acclimatized at our animal facility for 1 week before they underwent stereotaxic surgery under Isoflurane anesthesia (5 % induction, 1.5-3 % maintenance; O_2_: 0.5-1.0 L/min; Atm.Air.: 0.5-0.0 L/min). Fur on the head was shaved and cleaned 3 times with 70 % ethanol and 0.5 % chlorhexidine in 85 % ethanol. Pre-operatively the animals were administered Buprenorphine (Bupaq; 0.3 mg/mL; 0.03 mg/kg), Carprofen (Norodyl/Rimadyl; 50 mg/mL; 5 mg/kg) and sterile saline (0.9 %; 5 mL/kg). A mixture of Lidocaine (5 mg/kg, 10 mg/mL) and Bupivacaine (1 mg/kg; 5 mg/mL) was injected subcutaneously on the scalp, 10min before incision. The rectal temperature was monitored and maintained at 37.5 °C on a heat pad (Harvard Apparatus Homeothermic Monitoring System). Heart rate (HR) and oxygen saturation SpO_2_ was monitored during surgery (Nonin 2500A VET) and used to adjust isoflurane level and (O_2_:Atm.Air)-ratio. Viral inoculation as well as chronic implantation of the optic fiber, electrodes and pedestal were carried out in the same surgical session to reduce the number of surgeries per animal. All stereotaxic coordinates were normalized to a Bregma-Lambda distance of 9.00mm to correct for the smaller brain size of the 4weeks old animals [30].

Craniotomies were made above M1 using the following stereotactic coordinates relative to bregma (Fig.2): Anterior-Posterior (AP): +1.0 mm; Medial-Lateral (ML): +2.5 mm (right M1) and −2.5 mm (left M1), and dura was punctured. In right M1, 1 μL of pAAV-CaMKIIa-hChR2(H134R)-EYFP (UNC Vector Core) was injected at 0.2 μL/min, using a Dorsal-Ventral (DV) penetration depth of −1.00 mm relative to dura to reach L2/3 of M1.

**Figure 2:**
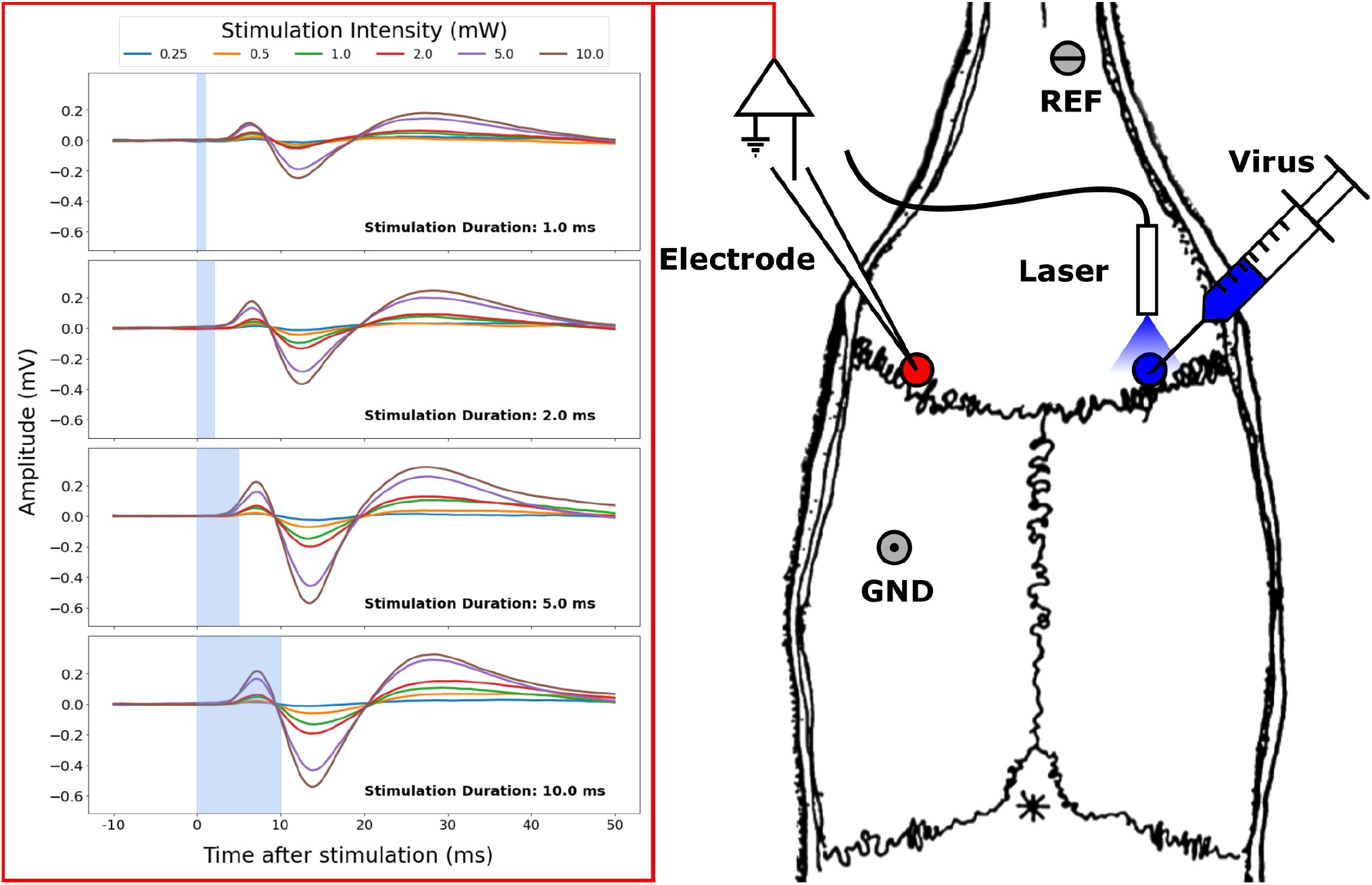
Transcallosal neural responses in left motor cortex (M1) evoked by contralateral optogenetic stimulation of right M1. The right panel shows the skull of a rat (adapted from [30]) indicating the stimulation site (right M1) and recording site (left M1). The viral optogenetic construct was injected into right M1 (blue circle), enabling subsequent optogenetic stimulation of cortical glutamatergic neurons in right M1 through an implanted optic fiber. Optogenetic excitation of transcallosal projection fibers to contralateral left M1 gave rise to transcallosal evoked potentials in left M1 (red circle). The left box depicts the mean transcranial evoked local field potential (LFP) recorded from one animal (rat27.2, 141 days old, 111days after surgery), showing an early positive component (P1) and a subsequent negative component (N1). The four panels depict responses evoked with optogenetic stimulation at four different stimulation durations out of seven durations in total. The different colors correspond to different stimulation intensities levels. REF: Location of the reference electrode. GND: Location of the ground electrode.

A custom-made and calibrated optic fiber implant (see previous section) was implanted at the same position (Fig.2). In left M1, a twisted electrode pair made of stainless steel (Plastics1; E363/3-2TW; two wires, Ø = 127 μm each) was implanted at a penetration depth of −1.00mm DV, relative to dura. Two screw electrodes (Plastics1; E363/20/2.4) were inserted into craniotomies for reference (AP: +8 mm; ML: +0.75 mm) and ground (AP: −3.00 mm; ML: −3.00 mm). All electrodes were connected to a 6-channel plastic pedestal connector (Plastics1; MS363). Optic fiber, depth and screw electrodes as well as the pedestal was cemented to the skull using dental acrylic cement (3M RelyX Unicem and GC Fuji Plus; or Panavia V5: Clearfil; Tooth Primer; Paste). Later animals had the implant embedded within a custom-made and 3d-printed implant protector (https://git.drcmr.dk/cskoven/lab) with an unmountable lid, in order to reduce damage to and dust on imaplants. The wound was sutured (Ethicon Vicryl, 5-0 Vicryl, FS3, 16 mm) and another volume of saline (5 mL/kg) was injected subcutaneously to accelerate rehydration. The animals were set to wake up on the heating pad with continuous flow of (O_2_:Atm.Air) in the gas mask, without isoflurane - and subsequently put back in a clean cage, with free access to water as well as solid and softened hydrated food.

### Postoperative treatment

Animals were allowed to recuperate one week in quarantine, with postoperative analgesic treatment of Carprofen (Norodyl; 50 mg/mL; 5 mg/kg) and antibiotic treatment of Baytril (50 mg/mL; 10 mg/kg), once daily for 5 days or as needed. Hereafter animals were returned to similar caging/housing, bedding and enrichment in the animal housing facility. Animals were not included in experiments until 4 weeks after surgery. The rat pedestal protector improved wound healing and allowed us to house two animals together hereafter.

### Measurement of the optogenetically evoked transcallosal LFPs

Optogenetic stimulation and electrophysiological measurements were carried out four to seventeen weeks after surgery

### Anesthesia

Animals were anesthetized with Isoflurane (5%; 1 L/min @ 50% O_2_ + 50% Atm.Air.), which was downregulated to 2-3 % for insertion of tail vein catheter. The catheter was flushed with heparinized sterile isotonic saline (Leo Pharma; 5000 IE/a.e./mL; 0.33 mL Hep. per 100 mL Saline) to avoid clotting. Heart rate (HR) and SpO_2_ was monitorized (Nonin 2500A VET) and logged/visualized with custom-made scripts (https://git.drcmr.dk/cskoven/nonin). A small bolus (“fast infusion”: ~0.1 mL/10s) of Dexmedetomidine (Dexdomitor; Orion Pharma; 0.1 mg/mL) was administered (Harvard Apparatus Pump 11, Pico Plus Elite) until a steep decrease in HR (~20 %; e.g., 375 bpm to 300 bpm within 15 seconds) was observed. Regular infusion was continued (0.05 mg/kg/hr for the first hour; 0.15 mg/kg/hr hereafter) in combination with a downregulated isoflurane level (0.5 %; 1 L/min) [31,32].

### Recording and stimulation setup

Animals were then placed in a Faraday cage (Campden Instruments; CI.80604E-SAC-H1). Body temperature was monitored and stabilized at ~37.5 °C on a heat pad (Harvard Apparatus Homeothermic Monitoring System) throughout the experiment. The fiber optic patch cable (L = 5 m, Ø = 50 μm, Thorlabs) was connected to the ceramic ferrule embedded in the cranial cement with a ceramic ferrule sleeve connector (Thorlabs ADAL1). The electrode pedestal was connected to a 4-channel differential bio-amplifier (Warner Electronics DP-304A) through a connector cable (Plastics1; 363-000) custom adapted to BNC-plugs. Gain was set to x1000 for the first animals (N=4), but later reduced to x100 in subsequent recordings to avoid clipping of the signal. Hardware filters were set to bandpass the signal between 0.1 Hz and 50 kHz. The amplified signal was recorded and digitized at 30 kHz on a Lenovo T460 laptop (Ubuntu Linux 18.04) using an Open-Ephys acquisition board and software (v.4.5.0) [33].

Once the animals were stable (HR, SpO_2_ ≥ 98 %, T = 37.5±1 °C) they were exposed to laser stimulation (447 nm) according to a pseudo-randomized parameter mapping paradigm as follows below. The laser and optic fiber patch cables were the same as used for previous calibration of the optic fiber implants. The stimulation variables of optogenetic stimulation were controlled by an open-source Pulse Generator (PulsePal rev.2, firmware v.2.0.1, Sanworks) [34] with custom-made python scripts (code available at https://git.drcmr.dk/cskoven/PulsePal).

### Optogenetic stimulation paradigm

We applied optogenetic stimulation to right M1 using different combinations of stimulation durations and stimulation intensity. We recorded the cortical response in left M1, evoked by optogenetic stimulation in right M1 and propagated directly through corpus callosum. In total, seven different stimulation durations (0.1 ms - 10.0 ms) and six different stimulation intensities (0.25 mW - 10.0 mW) were combined, resulting in 42 duration-intensity settings. For each duration-intensity combination, we applied 150 stimuli, recording a total of 6300 optogenetically evoked transcranial LFPs in the left M1. After the initial four experiments, we made some adjustments to improve signal-to-noise ratio. In the first four animals, we tested each of the six levels of stimulation intensities (0.25; 0.50; 1.00; 2.00; 5.00; 10.00 mW) in one block (gain: x1000). At each stimulation intensity, we randomly intermingled blocks of 50 trials at three different stimulus durations. Hence, stimulations at a given stimulation intensity were carried out in one train and were thus placed relatively fixed in the anesthetic paradigm. To spread out the various stimulation duration–intensity combinations throughout the experiment, we randomly intermingled blocks of multiple combination in all subsequent experiments. For each stimulation intensity, three trains were carried out having 2 blocks of 25 trials for each of the seven different stimulation durations, with these blocks being randomly intermingled (gain: x100). Initially, trial duration was fixed to 1000 ms which led to an accumulation of 50 Hz line noise during blocks of measurements. We therefore increased the trial duration slightly (to π/3 ~ 1.0472 s) to secure a trial-by-trial shift in phase for the underlying 50 Hz line noise in later experiments. In summary, our adjustments reduced the effects of line noise and potential impact of physiological variables as a function of the anesthetic depth. Since overall stimulation conditions were kept constant, data acquired before and after these adjustments were pooled. The change in signal amplification was handled during signal processing. A few trials (< 8 trials per condition) had to be discarded due to technical reasons.

### Data analysis

When exposing the right M1 to laser stimulation, the typical transcallosal response evoked in left M1 entailed a small positive peak (P1) and a subsequent and more prominent negative response (N1), corresponding to response pattern “1” in Figure 1b. In some animals, the P1 peak was absent or too small for automatic detection, corresponding to response pattern “2” in Figure 1b. Since the N1 peak was robustly detected in almost all animals, our analyses focused mainly on the N1. Data analysis was performed using custom-made scripts programmed in Python (https://git.drcmr.dk/cskoven/elphys) and open-source reading tools for electrophysiological data (https://github.com/open-ephys/analysistools). Preprocessing included notch filtering at 50 Hz and 2^nd^ order Butterworth band-pass filtering between 3 Hz and 300 Hz. Sessions recorded with a gain factor of x100 instead of x1000, was multiplied by a factor 10 as compensation. Single trials included all data points recorded 900 ms before and after the onset of optogenetic stimulation. Trials were baseline corrected using the mean value of the data points in the 10 ms period before laser stimulation onset. Trials were grouped according to duration-intensity condition (42 conditions: 7 stimulation durations and 6 stimulation intensities).

### Peak detection

For each condition, trials were averaged. The first positive (P1) and negative peak (N1) were automatically detected, if their amplitude extended beyond the threshold of 1 standard deviation based on the 100 ms preceding the stimulus onset. Peak detection was restricted to a time window of 1-9ms after stimulation onset for P1 and 5-20 ms for N1. Automatic peak detections were verified by CSS, to eliminate possible spurious detections. Peak onset was interpolated [35,36], and defined by the intersection of a first order regression line on the peak slope, between 45 % and 55 % of the peak maximum, and the baseline. We created color-coded group maps to describe the detectability, amplitude, latency and variability of the optogenetically evoked transcranial response in M1.

## Results

Measurements of transcallosal LFPs were performed in 17 animals four weeks after surgery. Eight animals were exposed to LFP measurements 9 to 17 weeks after surgery, including four animals who had been included in the early measurement. Five of the early and two of the eight late postoperative recordings had to be discarded due to noisy data or failure to elicit a transcallosal response with optogenetic stimulation. When showing group results, each animal is only represented once, even if two successful recording sessions were available. We thus report experimental data acquired in 15 sessions, nine sessions at young age and six sessions at older age.

We first assessed at which stimulation duration-intensity combination optogenetic stimulation in right M1 was able to evoke a consistent transcallosal LFP in the left M1 (Fig.3). The ability to reliably evoke a transcallosal response gradually increased with stimulation duration and intensity with the two variables having a synergistic effect. High stimulation intensity (5 or 10 mW) reliably evoked a transcallosal N1-response already at very short stimulus durations of 0.5 or 1 ms in all 15 recording sessions (Fig.3b,c). In contrast, a reliable P1 response could only be evoked in 11 of the 15 recording sessions (Fig.3a,d) even at stimulus combinations with high stimulation intensity and long stimulation durations. To systematically characterize the stimulation-response relationship between right M1 stimulation and transcallosal evoked respond in left M1, we focus in the following on the N1 peak as the output metric of interest.

**Figure 3:**
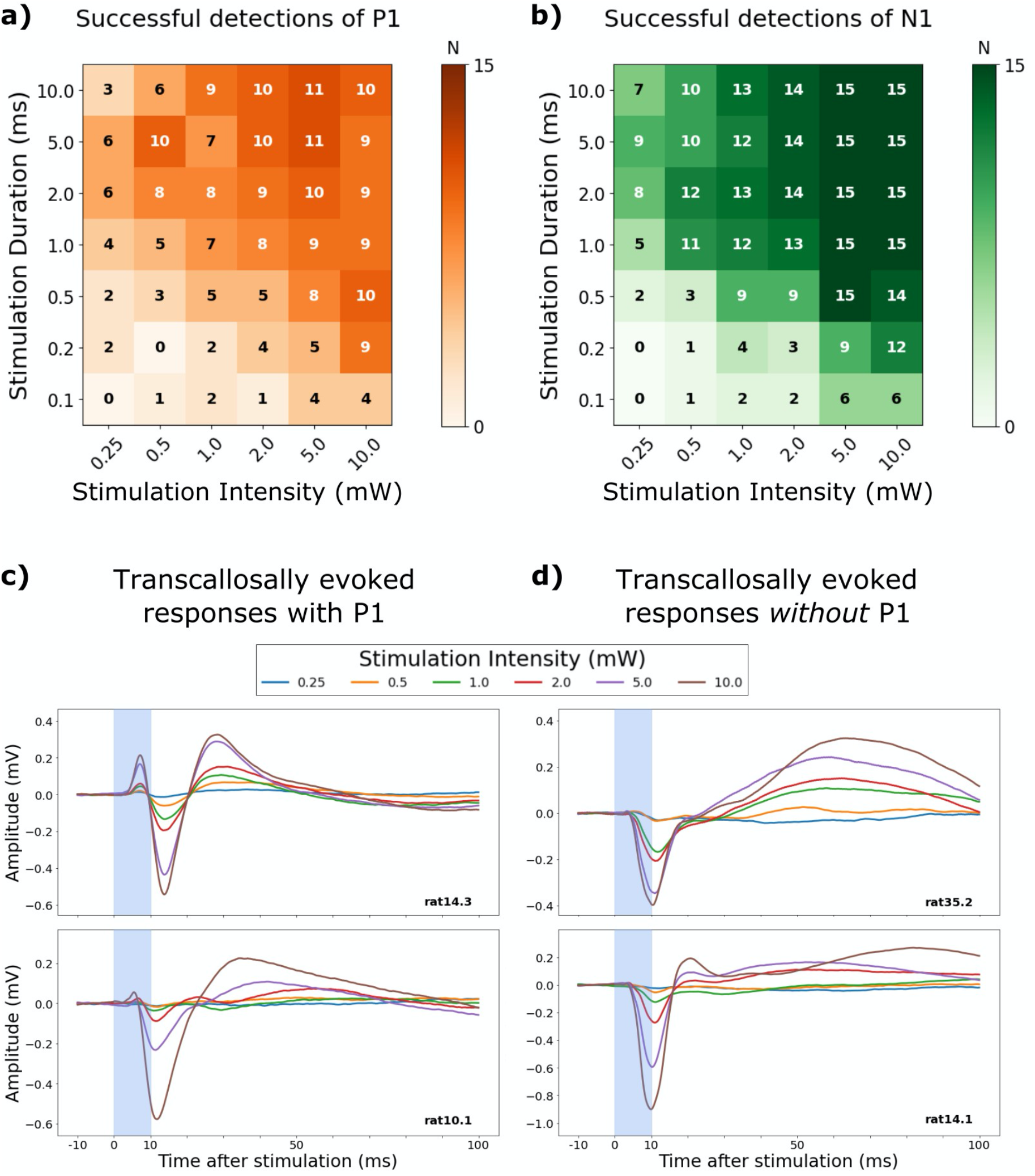
Detectability of a transcallosal P1- or N1-peak response depending on stimulation intensity and duration. **a,b)** The upper panels show color coded grids indicating the relative frequency of successful P1 peak detection (orange) or N1 peak detection (green) in left M1 after optogenetic stimulation of right M1. Each grid cell corresponds to one of the 42 combinations of stimulation intensity (x-axis) and duration (y-axis). The numbers plotted inside each cell of the grid indicate the absolute number of animals showing a detectable peak. The lower panels depict averaged samples traces of the transcallosal LFP of four animals. Two animals show a P1-peak **c)**, while a P1 peak is absent in the other two animals **d)**. Stimulation duration was fixed (10 ms). Stimulation intensities are indicated by color coding.

We further examined the response properties of the N1-peak in those stimulation conditions in which optogenetic stimulation of right M1 elicited a robust LFP in left M1 (Fig.4). N1-peak responses became less variable, as indexed by a lower coefficient of variation (CoV), when using a high stimulation intensity level and long stimulation duration (Fig.4a). This is underlined by the number of animal sessions with a CoV below a threshold of 1.0 for each individual condition (Fig.4b) The latencies of the N1 peak became gradually shorter with higher stimulation intensity, but a longer duration of the laser stimulus did not result in shorter latencies of the N1 peak (Fig.4c). This was most likely due to an emergence of a P1 peak that may result in a later peak of the N1 component.

**Figure 4:**
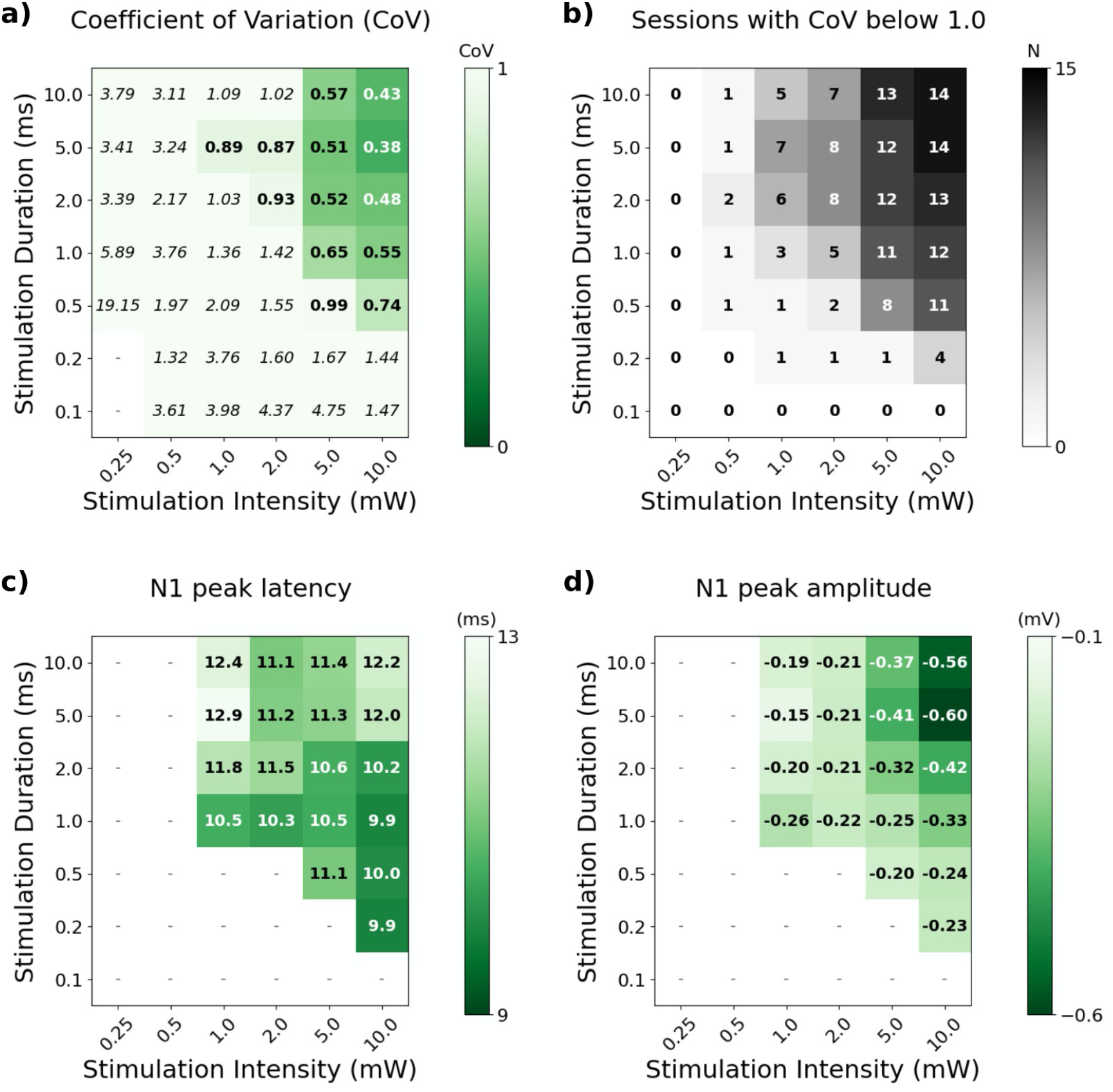
Stimulus-response characteristics of the N1-component of the transcallosal evoked LFP in left M1. **a)** Median of robustness of the N1 response across animals, determined as coefficient of variation (CoV; SD/mean) of peak N1 amplitude for individual recorded trials. We selected a CoV-threshold for robustness of the N1 peak of 1.0 (values below threshold are marked in bold). **b)** Number of animal sessions with robust conditions (CoV < 1.0). Median values for panel **c + d** are only shown where there is ≥ 3 animal sessions for a given condition. **c)** Median N1 peak latency (ms) after stimulus onset of robust conditions (CoV < 1.0) with N ≥ 3 animal sessions. **d)** Median N1 peak amplitude (mV) of robust conditions (CoV < 1.0) with N≥ 3 animal sessions.

Regarding response magnitude, transcallosal evoked response gradually increased with the “dose” of optogenetic stimulation (Fig.4d). When considering only those stimulation conditions that produced a robust transcallosal response, our measurements revealed a steady increase of N1 peak amplitude with stimulus intensity for stimulus durations between 2 and 10 ms (Fig.4d). Likewise, N1 peak amplitude gradually increased with stimulus duration, but only at relatively high stimulus intensities of 5 or 10 mW (Fig.4d).

Since the optic fiber and electrodes were chronically implanted, we were able to successfully elicit and record transcallosal cortical responses in two animals, approximately three months after the first parameter mapping experiment (Fig.5). In both sessions, we found a comparable increase in N1 peak amplitude with stimulation intensities. While there was a slight delay in N1 peak latency in the first rat (rat27.2), N1 latency did not change in the other (rat27.3). Contrary to its first session, a minuscule P1 was visible in the second session for rat27.2.

**Figure 5:**
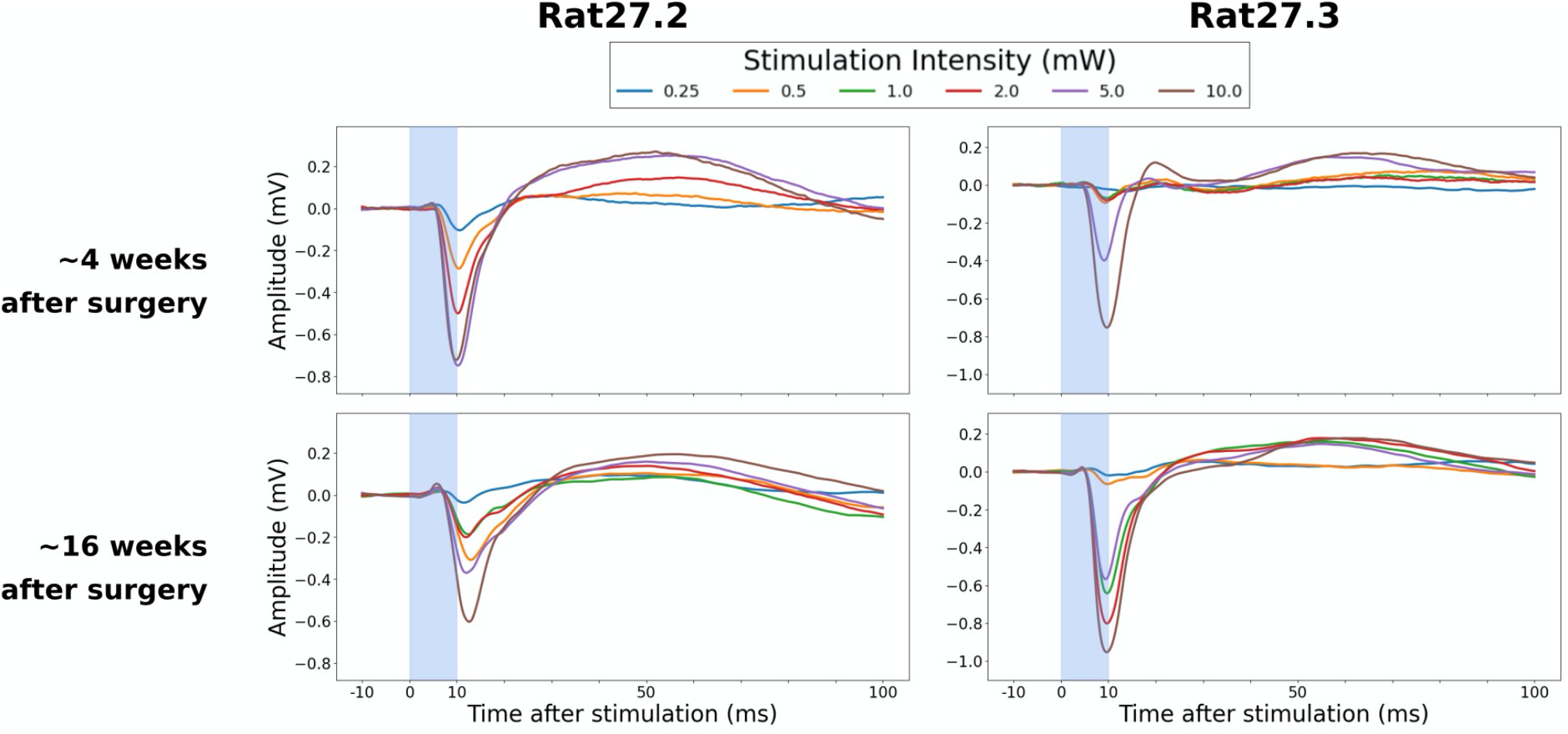
Repeated recordings of transcallosal LFPs in the left M1 in two rats. Colors correspond to the different stimulation intensities. Stimulus duration was 10ms (illustrated by a light blue rectangle overlay).

## Discussion

Recording the LFP potentials in rat M1, we were able to characterize the transcallosal cortical response elicited by optogenetic stimulation of glutamatergic neurons in the homologous contralateral M1. Systematic stimulation-response mapping revealed a gradual increase in the transcallosal M1 response, when increasing the duration and intensity of laser stimulation, - both individually and in combination. Conversely, the onset of the transcallosal M1 response did not show a gradual decrease in latency with increasing stimulus duration or intensity. In the discussion, we first compare our results with recent studies applying optogenetic stimulation to the transcallosal pathway and with classical electrical stimulation studies of the same pathway. We will then discuss the stimulus-response relationship, characterizing the relationship between the intensity and duration of optogenetic stimulation and the size and latency of the transcallosally evoked response. Finally, we will discuss the potential and limitations of our experimental approach and how it may be useful in future studies on the function and plasticity of transcallosal M1-M1 connections.

### Intracortical response evoked by transcallosal optogenetic stimulation

The cortical LFP responses elicited by optogenetic stimulation of glutamatergic transcallosal projections are in good agreement with previous optogenetic studies in rodents [25,27,37]. Using transgene rats with ubiquitous ChR2 expression, Saiki et al. [25] performed cortical optogenetic stimulation of M1 and recorded transcallosally evoked single-unit activity in contralateral M1 at a mean latency of 5.2 ms.

Transcallosal interhemispheric signal conduction has also been probed with optogenetic stimulation of the somatosensory [26] and barrel cortex [27,37]. Iordanova et al. [26] targeted specifically the glutamatergic neurons in the right somatosensory cortex and recorded the optogenetically evoked transcallosal response using silicon probes with linear electrode arrangement. This experimental set-up allowed them to compute the current source density (mV/mm^2^) throughout the cortical depth as a function of time after stimulus onset. Depth profiling of the LFP response showed that the polarity of the first LFP component flips with increasing depth of recording with an initial P1 peak emerging at deeper intracortical recording sites [26]. Chen et al. [27] recorded the transcallosally evoked responses after optogenetic stimulation of the somatosensory cortex, reporting a transcallosal response latency of approximately 8.1 ms. Böhm et al [37] performed epicranial optogenetic stimulation above the barrel cortex in mice, and recorded epicranial responses in multiple areas. They reported similar evoked potentials with delays and amplitudes depending on the recording location, illustrating how the transcallosally evoked response spreads among cortical areas.

Electric stimulation in cats, monkeys and rats consistently showed interhemispheric responses that closely resembled the transcallosal responses evoked with cell-type specific optogenetic stimulation of glutamatergic projections in the present study [5,6,38–40]. It has been shown in rat, cat and mouse that some GABAergic neurons also make transcallosal long-range connections with the contralateral cortex [41,42]. Electrical stimulation lacks cell-type specificity, and this should excite both glutamatergic and GABAergic transcallosal axons. The close resemblance of transcallosal potentials evoked by cell-type specific optogenetic stimulation and non-selective electrical stimulation suggests that the majority of transcallosal projections are indeed glutamatergic.

In our study, the most robust optogenetically evoked response was a negative peak (N1) in contralateral M1. In conditions in which optogenetic stimulation elicited a reliable N1 response, median N1 peak latencies ranged between 9.9 and 12.9 ms without showing consistent shortening of peak latency with increasing stimulation intensity or duration. A preceding positive peak (P1) was often detected, but only in approximately two thirds of the recording sessions. This variability can be attributed to differences in electrode locations with respect to the depth of the cortical column, giving rise to different configurations of the recorded local field potentials [26]. Our experimental approach was suited to reveal the magnitude of the transcallosally evoked cortical response, but the recording procedures were not geared to identify which cortical neuronal populations were primarily responsive to transcallosal optogenetic stimulation. Since the transcallosally evoked potential (N1 or P1-N1 response) reflects the summated activity of neuronal populations located close to the recording electrode [35,43], we cannot infer which cell types or microcircuits were preferentially activated by the transcallosal glutamatergic input. Considering the existing literature [23,42,44], we hypothesize that the bulk of the transcallosally evoked intracortical potential reflects transsynaptically evoked intracortical inhibition and its secondary impact on intracortical neuronal activity. Yet, more sophisticated experimental approaches would be required, offering cell-type or layer-specific activity readouts preferably combined with pharmacological manipulations to investigate the neuronal correlates of the transcallosally evoked potential at the intracortical microcircuit level. In addition, more elaborate stimulation protocols such as paired-pulse or burst stimulation may be suited to reveal the neuronal dynamics of optogenetically evoked, transcallosal M1-to-M1 interactions.

### Systematic evaluation of the optogenetic input parameters

The present study significantly extends previous work on optogenetic stimulation of transcallosal projections in rodents. We explored the impact of a various combinations of stimulus intensities and durations on the magnitude of the transcallosally evoked cortical potential, covering the most commonly used stimulation variables [20,28,45–51]. We show that the probability to reliably evoke a transcallosal cortical response gradually increased with the intensity and duration of the optogenetic stimulus. The transcallosally evoked cortical potential also became more stable as reflected by a gradual reduction of between-trial variability. This was the case for the N1 and P1 response. The increase in response stability and consistency was paralleled by an increase in magnitude of the transcallosally evoked potential at higher stimulus intensities and longer stimulus durations. This stimulus-response pattern mirrors the pattern known from similar to classical electrical stimulation [6], where the cortical neuronal response is determined by the relationship between the induced electrical current and the electrical properties of the neural tissue [6,52–54]. In the case of optogenetic stimulation, the efficacy to trigger axonal firing depends on how effectively the applied laser light activates light-sensitive ion channels in the neurons expressing channelrhodopsins [55–58]. The longer the exposure to laser light and the higher the laser power, the larger the regionally induced response in sensitive neurons. Taken together, our results show that it is possible to exploit the biophysics of optogenetic stimulation to probe the “gain function” of transcallosal M1-to-M1 glutamatergic projections in the intact rat brain. When interpreting the doseresponse relationship as revealed by “optogenetic dose” and evoked transcallosal response magnitude, one needs to bear in mind that variations in dose will alter the area of effective stimulation. Optogenetic stimulation at higher amplitude and longer duration may not only more effectively activate neurons that are located in the “hotspot” region of stimulation but also more effectively excite neurons in the brain tissue surrounding the hotspot, reducing anatomical specificity of the optogenetic intervention.

In our LFP measurements, response magnitude still continued to increase with increasing intensity and stimulation, even at high laser power levels and long stimulus durations (Fig.4d). However, we rather expected a sigmoid dose-response function with N1 peak amplitude reaching a plateau at high stimulus durations and intensities. We therefore conclude that the intensity-duration combinations used in this experiment did not cover the entire range of possible dose-response relationships. However, high stimulation intensity could cause heating of the tissue, which introduces spurious findings in functional MRI [48], may result in tissue damage [59–63], and may illuminate an unreasonably large tissue volume, hampering spatial specificity [25]. Indeed, care should be taken to restrict the stimulation volume to the transfected neuron population in the brain region of interest [20]. Hence, it may be advisable to restrict variations in stimulus intensity and duration to the low-to-moderate dose range, when studying dose-response relationships with optogenetic stimulation.

Since our examination also covered the lower range of intensity-duration combinations, the measurements also give indications regarding the minimal amplitude and duration needed to obtain a reliable transcallosal responses in studies that are not directly examining the dose-response relationship (Fig.4). From our data, one can conclude that intensities of 5-10 mW combined with durations of 1-10 ms are sufficient in most animals to elicit a robust transcallosal response.

In electrical stimulation studies, a short stimulus duration is usually desired, to minimize the duration of the electrical artifact and to better depict the direct result of the stimulation. The former concern is less of an issue in optogenetic studies, as there are no immediate stimulation artifacts, although some report photo-electric artifacts when using optrodes [59,64]. But an unnecessarily long stimulation duration may complicate the physiological interpretation of the optogenetically evoked cortical response. Interestingly, median latency of the transcallosally evoked N1 peak did not follow the same gradually increasing dose-response pattern as we found for N1 peak amplitude. Median N1 peak latency was shortest at the highest stimulus intensity (10 mW), but only when using relatively short stimulus durations, ranging between 0.2 and 2 ms (Fig.4c). At higher stimulation durations, an additional P1-peak emerged in several animals. The emergence of an additional P1-peak prolonged N1-peak latency in these animals, resulting in an apparent “delay” in median N1-peak latency at the group level. Therefore, combining a relatively high stimulus intensity with a relatively short stimulus duration may be advantageous, when estimating the peak latency of the transcallosal N1 response.

### Implications and limitations

Our LFP results highlight the importance to characterize the dose-response relationship in optogenetic stimulation studies. Detailed knowledge about the dose-response properties is critical to the selection of the optimal stimulation parameters and to understanding the evoked neuronal response. A recent study measured locally evoked spiking and LFP activity at the site of cortical optogenetic stimulation in awake, alert non-human primates confirms our conclusion [28]. In that study, the selection of stimulation variables determined the stimulation-evoked, excitatory-inhibitory responses, highlighting the role of in-depth understanding of dose-response relationships in optogenetic stimulation studies.

Our results also form a solid basis for future studies on the functional integrity of the motor transcallosal fibers in rodents. Since our approach allows to reliably investigate functional aspects of a specific cortico-cortical interhemispheric pathway, a next obvious step will be to add non-invasive microstructural mapping techniques to relate the dose-response relationships at the functional level with the microstructural properties of the transcallosal fiber tract, including *post mortem* histological validation [65,66]. Further, this preclinical platform can be used translationally to assess animal disease models (e.g. multiple sclerosis, stroke, trauma) that affect the transcallosal M1-M1 pathway. This will open up novel possibilities to study the function-structure relationship of the transcallosal motor pathway and how it is affected by disease.

In two rats we were able to demonstrate the feasibility of longitudinal recordings, showing that robust LFPs could be obtained in two measurements made approximately eight weeks apart. The possibility to repeatedly probe directional functional connectivity between the right and left M1 opens up for longitudinal investigations of e.g., how transcallosal motor pathways are functionally shaped by maturation, aging, specific brain diseases, or experimental manipulations (i.e., training, pharmacological challenges).

As pointed out in the introduction, dual-site TMS, targeting the left and right M1, is a well-established tool in humans to study interhemispheric interactions between the motor cortices [7,8,15,67–69]. Using a conditioning-test paradigm, the conditioning pulse given over one M1 produces a prominent inhibition of cortical excitability in the contralateral M1. This transcallosal cortico-cortical inhibition already emerges, when the conditioning stimulus is given 7-12 ms before the contralateral test pulse [7]. Of note, this inhibitory period is often preceded by a short period of interhemispheric facilitation which occurs at an inter-stimulus interval of 4-5 ms [69], suggesting that the conditioning TMS pulse transcallosally triggers a mixture of early excitation followed by inhibition in the conditioned M1. This dual-site M1-M1 stimulation protocol has been intensively used in clinical populations, including patients with multiple sclerosis or motor stroke [14,15,70]. Interventional repetitive TMS of human M1 has also been used in humans to alter the strength of transcallosal motor interactions, especially in patients with motor stroke to mitigate a functional imbalance between the lesioned and non-lesioned hemisphere [71,72]. Interventional TMS approaches to attenuate interhemispheric imbalance after unilateral cortical injury have also been explored in the rat using implanted electrodes [73]. Here, our optogenetic stimulation approach in rats offers the possibility for more specific targeting of neural elements underlying interhemispheric M1-M1 interactions and for identifying the most relevant stimulation variables. This knowledge may be used to inform the optimization of interventional TMS approaches in humans.

The latter could be introduced by for instance long-term exposure or by LTD- or LTP-inducing paradigms (1 Hz vs. 10 Hz [74]). Finally, this approach adds insight to neurobiological questions also being probed in humans, where the level of neural detail is restricted to what can be obtained non-invasively.

### Methodological considerations

The study has some methodological limitations. Electrophysiological recordings and analyses focused on single-electrode data and population responses as reflected by the LFP. The use of multi-electrode arrays covering different depth levels of the cortex and analysis of multi-unit activity would have yielded a more complete picture of the transcallosally evoked cortical response pattern with respect to cortical layers [75]. Our approach also was not able to disentangle the contribution of various cell types, e.g., pyramidal cells and inhibitory interneurons, to the transcallosally evoked LFP. Pharmacological manipulations or the use of cell-specific readouts of functional activity may be used in future studies to dissect the various components of the transcallosal response at the micro-circuit level. A similar consideration applies to optogenetic stimulation. The use of more cell-type specific viral vectors might help to reveal which cortical cell types are the main contributors to the transcallosal interaction and how they can be most effectively stimulated and modulated with interventional neurostimulation.

All recordings were performed under general anesthesia to have a setup where the brain state is somewhat stable, but this limits the translation of the results to the physiological states present in normal behaving wake animals. Our anesthesia protocol was optimized for future concurrent use with MRI [32,76] which will enable us to investigate structural and functional transcallosal connectivity using several modalities.

## Conclusion

By systematically mapping the relationship between “stimulation dose” and “response magnitude”, we were able to characterize the gain function of transcallosal M1-to-M1 glutamatergic connections in intact rats, using a set-up that is suited for long-term *in vivo* stimulation and recordings. Combining unilateral optogenetic stimulation of glutamatergic neurons with intracortical recordings of the transcallosally evoked response has substantial translational potential to foster a better understanding of functional M1-M1 transcallosal interactions and to inform interventions that use transcranial cortex stimulation to modify the balance in transcallosal M1-M1 interaction.

The observed dose-response profiles of the N1 peak in terms of variability, magnitude and latency suggest that one should aim at finding the optimal trade-off between stimulation duration and intensity in optogenetic studies. Although evoked responses become more robust at higher stimulation intensities and longer stimulation durations, other intensity-duration combinations may be preferable depending on the research question. In many cases, the intensity-duration combination needs to be sufficiently strong to elicit a robust transcallosal response, but not too strong to secure sufficient sensitivity towards dynamic changes in transcallosal connectivity that are experimentally induced or arise across the lifespan.

## Acknowledgements

We wish to thank our local Laboratory Technician, Sascha Gude, for her immense help with animal caretaking and assisting in experiments, throughout the project period.

We also wish to thank Søren Ellegaard for helping with the electrophysiological setup.

